# Cross-modal and non-monotonic representations of statistical regularity are encoded in local neural response patterns

**DOI:** 10.1101/243550

**Authors:** Samuel A. Nastase, Ben Davis, Uri Hasson

## Abstract

Current neurobiological models assign a central role to predictive processes calibrated to environmental statistics. Neuroimaging studies examining the encoding of stimulus uncertainty have relied almost exclusively on manipulations in which stimuli were presented in a single sensory modality, and further assumed that neural responses vary monotonically with uncertainty. This has left a gap in theoretical development with respect to two core issues: i) are there cross-modal brain systems that encode input uncertainty in way that generalizes across sensory modalities, and ii) are there brain systems that track input uncertainty in a non-monotonic fashion? We used multivariate pattern analysis to address these two issues using auditory, visual and audiovisual inputs. We found signatures of cross-modal encoding in frontoparietal, orbitofrontal, and association cortices using a searchlight cross-classification analysis where classifiers trained to discriminate levels of uncertainty in one modality were tested in another modality. Additionally, we found widespread systems encoding uncertainty non-monotonically using classifiers trained to discriminate intermediate levels of uncertainty from both the highest and lowest uncertainty levels. These findings comprise the first comprehensive report of cross-modal and non-monotonic neural sensitivity to statistical regularities in the environment, and suggest that conventional paradigms testing for monotonic responses to uncertainty in a single sensory modality may have limited generalizability.

## 1. Introduction

Currently, one of the dominant frameworks for understanding brain function couches perception in terms of learning-based predictive processes, which operate by integrating information over multiple temporal scales (e.g., Bornstein & Daw, 2012; Clark, 2013; Friston, 2010). This is a foundational premise in computational and cognitive approaches to economic decision-making, language processing, statistical learning, and low-level sensory processing. These theoretical developments have been accompanied by a rich body of experimental data addressing the neurobiological basis of predictive processing, and in particular, brain systems that encode temporally-unfolding statistical structure in the environment (see Hasson, 2017, for recent review).

There are, however, two substantial limitations to our understanding of the neurobiological systems encoding environmental statistics. First, almost all empirical studies probing the neural systems supporting predictive processing assume that these systems track statistical regularities monotonically; i.e., that the relevant neural systems are ones in which activity increases or decreases monotonically with statistical regularity, predictability, or uncertainty. This assumption is deceptively intuitive and is sufficiently ingrained in neurobiological experiments that it is rarely stated explicitly. Examples include neuroimaging studies that test for linear relations between statistical regularities in stimulus series and response magnitudes (e.g., Bischoff-Grethe, Proper, Mao, Daniels, & Berns, 2000; Harrison, Duggins, & Friston, 2006; Huettel, Mack, & McCarthy, 2002; Strange, Duggins, Penny, Dolan, & Friston, 2005), or studies that contrast structured and random inputs sequences (e.g., Cunillera et al., 2009; McNealy, Mazziotta, & Dapretto, 2006).

That said, there are a few recent exceptions to this assumption. Kidd et al. (2012) found that infants, when presented with sequences of events varying in their predictability (surprisal), were less likely to look away from intermediately surprising events than when events were too predictable or too surprising. This suggests that stimuli of intermediate predictability may receive privileged neural processing with respect to random or highly structured inputs. Along these lines, Nastase et al. (2015) found that whole-brain connectivity between the anterior cingulate cortex and several brain regions tracked statistical regularities in auditory stimuli non-monotonically (i.e., via a quadratic trend). Non-monotonic responses to regularity are compatible with several types of operations (Hasson et al., 2017; Nastase et al., 2015). For example, neural systems modeling the environmental generators of sensory inputs may be maximally engaged by moderately structured inputs where model complexity is highest; additionally, systems supporting exploratory behavior or encoding particular information-theoretic metrics, such as predictive information rate (Abdallah & Plumbley, 2009), may be maximally engaged by inputs of intermediate regularity, while not differentiating highly random and highly structured inputs. Identifying brain systems that respond non-monotonically to uncertainty, particularly ones that do so in a supra-modal manner, would expose a novel but unappreciated aspect of neural coding of input statistics, which cannot be explained by low-level mechanisms such as the construction of prediction, generation of prediction errors, or any other computational account in which responses scale with uncertainty.

A second, related limitation is that very few studies have directly investigated whether there exist neural systems that are sensitive to input statistics in more than one modality. This is one of the core questions in functional theories of statistical learning (for review, see, e.g., Frost, Armstrong, Siegelman, & Christiansen, 2015) but has seldom been addressed from a neurobiological perspective. Our prior work examining this issue (Nastase et al., 2014) failed to identify areas sensitive to regularity in both auditory and visual inputs (and was agnostic to the issue of linear or non-monotonic trends). Other work in which participants were instructed to predict the final elements of series varying in predictability reported adjacent (Schubotz & von Cramon, 2002) or overlapping responses (Schubotz & von Cramon, 2004) in left ventral premotor cortex for different modalities. More recent work (Meyniel and Dehaene, 2017) has implicated a more widespread network including precentral, intraparietal, and superior temporal cortices in tracking transition probabilities in auditory and visual series (though no formal conjunction test was performed). Nonetheless, overlapping activations (i.e., conjunction maps) provide limited evidence for cross-modal representation of uncertainty, because the finer-grained organization of neural activity may differ across modalities.

The current study was designed to address these limitations by (*a*) systematically probing for both monotonic and non-monotonic neural responses to statistical regularities in auditory, visual, and audiovisual stimuli, and (*b*) determining to what extent these responses are modality-independent. Participants were presented with brief ~10 s auditory, visual, and audiovisual series varying across four levels of entropy (i.e., uncertainty, inversely related to regularity) while performing an orthogonal cover task. The same statistical constraints were used to generate auditory series consisting of pure tones, visual series consisting of simple colored shapes, or audiovisual series where each token was a unique tone/shape combination (thus the uncertainty and structure of the audiovisual series was identical to that of the auditory-only and visual-only series). We used multivariate pattern analysis to localize response patterns that differentiated series with varying uncertainty in either a monotonic or quadratic (i.e., non-monotonic) fashion for auditory, visual, and audiovisual series. We also used multivariate cross-classification to identify neural systems encoding uncertainty in a modality-general fashion by training a classifier to discriminate levels of uncertainty in one modality (e.g., visual series) and then testing it on another modality (e.g., auditory series). This procedure explicitly tests for systems coding for statistical regularities in the environment at a level of abstraction that supersedes the sensory features of the stimuli.

In general, we expected different brain systems to track uncertainty within auditory and visual streams, consistent with emerging views that different neural systems encode sequential structure or environmental regularities in different modalities (for recent reviews; see Armstrong, Siegelman, & Christiansen, 2015; Dehaene, Meyniel, Wacongne, Wang, & Pallier, 2015; Frost et al., 2015; Hasson, 2017; Milne, Wilson, & Christiansen, 2018). We further expected that the systems implicated in tracking the level of uncertainty in audiovisual stimuli would diverge from those tracking uncertainty for unisensory stimuli, as our recent work (Andric, Davis, & Hasson, 2017) indicates that audiovisual inputs trigger unique computations related to uncertainty.

## 2. Methods

### 2.1. Participants

Twenty-five right-handed adults (Mean Age = 26.1 ±4.74 SD; 11 female) participated in the study, which was conducted at the University of Trento, Italy. They were recruited from the local student population, provided informed consent, and were reimbursed at a rate of 10 Euro per hour. Partic-ipants reported no history of psychiatric illness, history of substance abuse, or hearing impairments, and underwent an interview with a board-certified medical doctor prior to scanning to evaluate other exclusion criteria. Data from one participant who had completed the study were not included due to excessive movement during the scanning session. The human research ethics committee of the University of Trento approved the study. The data collected here have not been reported in any other study

### 2.2. Design, materials, and procedure

Stimulus events consisted of brief auditory (A), visual (V), or audiovisual (AV) series. Each series consisted of 32 items presented within 9.6 s at a rate of 3.3 Hz. These 32 items consisted of a repeated sampling of four tokens whose presentation order was determined by a first-order Markov process. For all modalities, each token was presented for 250 ms, followed by a 50 ms pause. For the auditory series, these tokens were four pure tones (262, 294, 330, 349 Hz; corresponding to middle C, D, E, and F in the Western major scale). Volume was manually adjusted for each participant until auditory stimuli were comfortably heard over scanner noise. For the visual series, the four tokens were four visual stimuli each identified by a unique combination of shape (circle, square, star, triangle), color (blue, green, red, yellow), and location (left, right, above, or below the fixation cross; e.g., ‘1’ = blue triangle presented above the fixation cross). Visual stimuli were presented at 2° visual angle from the fixation cross so that they could be observed without eye movement. The AV series consisted of yoked auditory and visual stimuli (fixed pairs) that were completely mutually informative such that within each AV series, any given tone was presented with only one visual stimulus. For each of the AV series, this produced an “alphabet” of only 4 possible states, analogous to the formal information content within the unisensory (A or V) series. The complete mutual information between auditory and visual streams in the AV condition reflected a single generating process, and consequently, tracking one stream provided complete information about the other. The specific instantiation of the one-to-one matching between a tone and a visual stimulus changed across the different AV series.

Series in the four conditions were generated using a first-order Markov process applied to four transition matrices with different levels of Markov entropy (Markov entropy = 0.81, 1.35, 1.56, 2.0; see Figure 1). We manipulated only these transition probabilities between tokens, while fixing the marginal frequencies across conditions at 25% per token; i.e., only Markov entropy was manipulated whereas Shannon entropy was fixed at 2 bits (a uniform distribution where each token is equally likely). We created the experimental series by repeatedly generating series from the Markov processes, evaluating those for transition constraints and marginal frequencies, and selecting only those series that exactly fit the generating process in terms of transition and marginal frequencies. Levels of the Markov entropy factor are referred to as levels 1, 2, 3, and 4 and indicate an increase in randomness; note that a positive linear relationship with entropy or uncertainty can also be described as a negative linear relationship with regularity, structure, or predictability.

**Figure 1:**
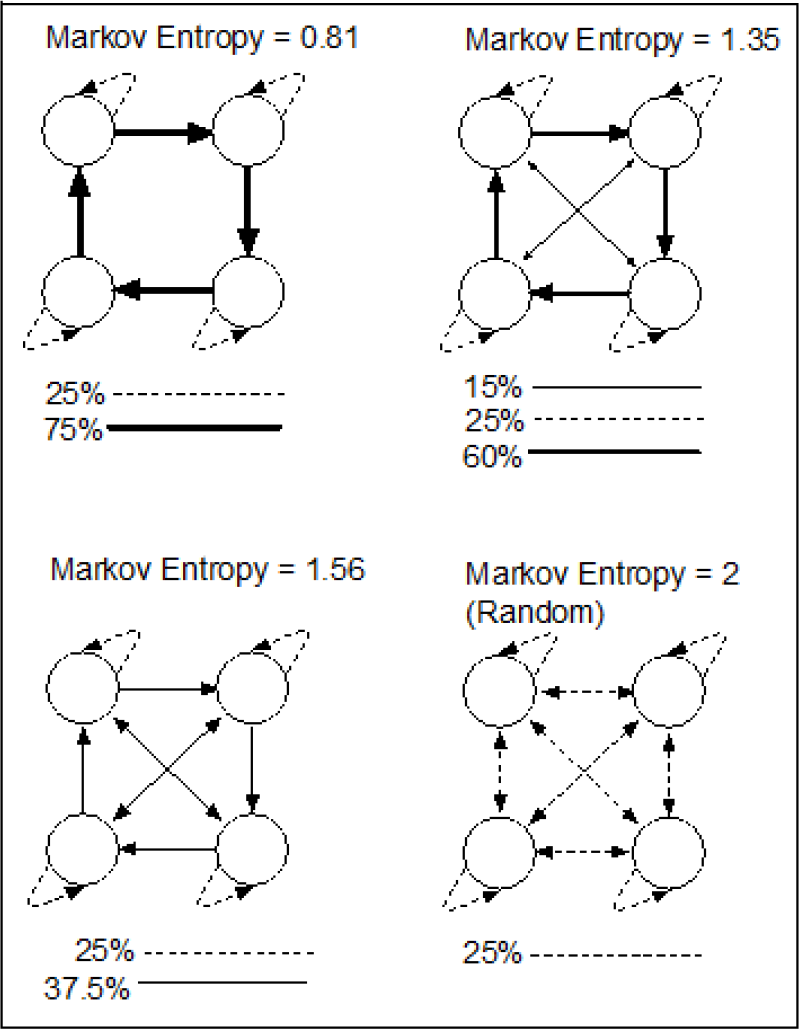
Transition graphs determining the uncertainty of stimulus series. In the auditory condition, each token corresponded to a tone. In the visual condition, each token corresponded to a shape of a particular color at one of four locations surrounding the central fixation cross. In the audiovisual condition, each token corresponded to a fixed tone-shape pair that remained unchanged within a series, but differed across series. The transition graphs correspond to entropy labels 1 (top left), 2 (top right), 3 (bottom left), and 4 (bottom right) in the text and figures.

There were 12 conditions in the factorial design corresponding to the fully crossed 4 entropy levels and 3 sensory modalities (A, V, AV). We used different series for each of these 12 conditions (i.e., 144 experimental series in total). These stimuli were presented over four experimental runs, with each run containing three series from each of the 12 conditions. Participants performed an orthogonal cover task in which they were instructed to monitor the fixation cross at the center of the display, and press a response key whenever the fixation cross began to rotate and alternate in color. These events served as catch trials, occurring six times during each run, and were unrelated to the entropy and modality manipulations. During each of the four experimental runs, performance was monitored online and not analyzed further; responses to catch trials were tracked by the experimenter and participants were provided feedback at the end of each run if a response was missed to encourage improved performance. In contrast to studies that encourage or require explicit prediction (e.g., Schubotz & von Cramon, 2002, 2004), we used a cover task to measure passive sensitivity to sensory regularities. Existing work suggests that explicit and implicit statistical learning tasks engage distinct, but partially overlapping neural systems (e.g., Aizenstein et al., 2004). The trial timing for each run was based on a rapid event-related fMRI protocol with jittered inter-stimulus intervals and an implicit baseline consisting of observation of the fixation cross. The presentation sequence was determined by the optseq utility (Dale, 1999), which generates a trial set optimized for this type of experimental design. Each run began with an 18.7 s rest interval to allow for signal stabilization.

### 2.3. fMRI acquisition

Images were acquired with a 4T MRI scanner (Bruker Medical, Ettlingen, Germany) using a birdcage-transmit, 8-channel receiver head coil (USA Instruments, Inc., OH, USA). Two T1-weighted 3D MPRAGE structural images were acquired (1 mm^3^ isotropic voxels, GRAPPA iPAT = 2, 5:36 min each). One was optimized for optimal contrast (MPRAGE_CNR) between gray and white matter tissue (TE/TR/TI/flip angle = 4.18 ms/2700 ms/1020 ms/7°) and the other was optimized for signal to noise ratio (MPRAGE_SNR) in gray and white matter tissue (TE/TR/TI/flip angle = 3.37 ms/2500 ms/1200 ms/12°; Papinutto & Jovicich, 2008). For functional MRI, single-shot EPI BOLD functional images were acquired using the point-spread-function distortion correction method (Zaitsev, Hennig, & Speck, 2004). Two hundred and eighty-five EPI volumes lasting 627 s in total were acquired during each of the four functional runs for 1,140 total volumes and 2,508 s total acquisition time (TR/TE = 2.2 s/33 ms, matrix 64 × 64, with 37 interleaved slices parallel to AC/PC, 3 mm^3^ isotropic voxels, slice skip factor = 15%, flip angle = 75.0°). Cardiac and respiratory measurements were not collected during fMRI acquisition.

### 2.4. Preprocessing

Preprocessing of fMRI data was carried out using FEAT (FMRI Expert Analysis Tool) Version 6.00, part of FSL (FMRIB’s Software Library; Jenkinson, Beckmann, Behrens, Woolrich, & Smith, 2012). The first six volumes of every fMRI run were discarded prior to analysis. The following preprocessing steps were then applied: motion correction using MCFLIRT (Jenkinson, Bannister, Brady, & Smith, 2002); slice-timing correction using Fourier-space time-series phase-shifting; non-brain removal using BET (Smith, 2002); spatial smoothing using a 5 mm FWHM Gaussian kernel; grand-mean intensity normalization of the entire 4D dataset by a single multiplicative factor; and high-pass temporal filtering (Gaussian-weighted least-squares straight line fitting, with sigma = 50.0 s).

In order to control for motion, confound matrixes were created using the dvars metric (Power, Barnes, Snyder, Schlaggar, & Petersen, 2012) using the fsl_motion_outliers tool. The dvars metric quantifies intensity differences between adjacent volumes after realignment (motion correction). Volumes that exceeded a boxplot cutoff threshold of 1.5 times the interquartile range were included in a confound matrix to be excluded in the first-level regression model by treating them as a regressor of no interest. This method is similar to excluding outlier time points from the regression model, but does not adversely affect temporal filtering or the autocorrelation structure.

### 2.5. Regression model

Single-participant analyses were conducted using FSLs FEAT (Jenkinson et al., 2012). A general linear model was constructed using FILM with local autocorrelation correction (Woolrich, Ripley, Brady, & Smith, 2001). The regression model included 12 regressors of interest: 4 (entropy levels) × 3 (sensory modality), where each 9.6 s event was modeled as a boxcar function convolved with a single-gamma hemodynamic response function. Note that we did not model or analyze the individual tokens (occurring at 3.3 Hz) comprising each event. Regressors of no interest included the catch trials eliciting button presses as well as a set of standard and extended motion parameters: 6 standard motion regressors, as well as their squares, temporal derivatives, squared derivatives, and the motion confound matrix determined using the dvars metric. We did not include a global signal covariate as stimulus series were brief (corresponding to approximately four functional volumes) and presented in pseudorandom order with jittered onsets, and because global signal regression may alter the inter-voxel correlation structure to which multivariate analysis are sensitive (Caballero-Gaudes & Reynolds, 2016).

### 2.6. Normalization

The structural images optimized for contrast-to-noise ratio (CNR) were preprocessed using the fsLanat tool according to the following steps: reorientation to MNI conventions (fslreorient2std); automatic cropping (robustfov); bias field correction (FAST); nonlinear normalization to a whole-brain MNI template (FNIRT); brain extraction based on the alignment to the MNI template, and segmentation according to tissue type and subcortical structures.

After estimating the first-level regression model, we aligned the statistical maps to MNI space in a single step by concatenating three transformation matrices resulting from the following three alignment stages. In a first step, each structural image was aligned to the first EPI volume in each run (i.e., the first of the six discarded volumes; the image with the strongest anatomical contrast) using a 3 degrees-of-freedom (translation-only) linear FLIRT alignment. In a second step, boundary-based registration (Greve & Fischl, 2009) was used to co-register the first EPI volume to the bias corrected, skull-stripped, and segmented structural image. In a third step the structural image was nonlinearly aligned to the MNI template using an initial 12 degrees-of-freedom linear registration step followed by nonlinear registration with a warp resolution of 10 mm.

### 2.7. Multivariate pattern analysis

#### 2.7.1. General approach and preparation for MVPA

We used multivariate pattern analysis to classify entropy conditions from distributed neural response patterns, with a focus on the issue of cross-modal classification (Kaplan, Man, & Greening, 2015; Kriegeskorte, 2011; Nastase, Halchenko, Davis, & Hasson, 2016). To localize brain areas that contained information about entropy, classification was performed using spherical volumetric searchlights (e.g., Kriegeskorte, Goebel, & Bandettini, 2006; Pereira, Mitchell, & Botvinick, 2009). Each searchlight had a 3-voxel radius (6 mm), and on average included 107 voxels (SD = 21 voxels).

We performed two types of classification where the classification targets were assigned so as to capture the two types of dissociations that were of theoretical interest: (*i*) classification of high versus low entropy conditions (approximating a “linear profile”); and (*ii*) classification of the two extreme (high, low) versus the two intermediate levels of entropy (“quadratic profile”). In this latter classification analysis, we assigned the label “extreme” to entropy levels 1 and 4, and the label “intermediate” to the entropy levels 2 and 3, and then proceeded with standard two-class classification. For the classifier to perform at 100% accuracy, it must, in left-out test data, classify both levels 1 and 4 as “extreme”, and levels 2 and 3 as “intermediate”.

First, we applied standard within-modality pattern classification to identify brain regions that discriminated levels of uncertainty in a linear or quadratic fashion. In this procedure, classifiers were trained and tested on response patterns *within* the same sensory modality (A, V, AV).

Second, to evaluate cross-modal sensitivity to entropy (i.e., information about entropy condition that generalizes across sensory modality), classifiers were trained on stimuli in the auditory modality and tested on stimuli in the visual modality (and vice versa) and the results averaged (as in, e.g., Man, Kaplan, Damasio, & Meyer, 2012; Oosterhof, Tipper, & Downing, 2012). Note that the audiovisual condition was not examined in the cross-classification scheme.

#### 2.7.2. Implementation of MVPA

We extracted regression coefficients from the first-level univariate general linear model, for each of the 12 conditions, and propagated those to a gray matter mask comprising the union of individual gray masks across participants (50% gray matter probability from FSL’s FAST) in MNI space. This gray matter mask contained 196,634 2 mm^3^ voxels after removing any voxels invariant across all samples and participants. Regression coefficients were averaged across the four runs within each participant prior to classification analysis to create a single map per condition, and then normalized (Z-scored) across features (voxels) within each searchlight (Misaki, Kim, Bandettini, & Kriegeskorte, 2010; Nastase et al., 2016). This normalization scheme ensures that the classifier cannot capitalize on regional-average differences in response magnitude (i.e., within a searchlight) between the different conditions.

Classification was performed using linear support vector machines (SVMs; Boser, Guyon, & Vapnik, 1992) with the soft-margin parameter C automatically scaled to the norm of the data. All classification analyses were performed using leave-one-participant-out cross-validation (e.g., Clithero, Smith, Carter, & Huettel, 2011; Mourao-Miranda, Bokde, Born, Hampel, & Stetter, 2005). That is, for each cross-validation fold, the decision boundary was constructed based on samples from 24 of the 25 participants, and tested on the left-out participant. This procedure was repeated until each participant served as the test participant, and the classification accuracies were averaged across cross-validation folds. It has been shown (e.g., Allefeld, Gorgen, & Haynes, 2016) that a leave-one-participant-out procedure more rigorously ensures that classification generalizes across participants than applying standard second-level statistical tests to classification accuracies that are based on leave-one-run-out cross-validation within participants.

In the cross-modal classification analysis, for each cross-validation fold the decision boundary was constructed based on samples from one sensory modality (e.g., auditory) in 24 participants, then tested on samples from the other sensory modality (e.g., visual) in the left-out participant. All multivariate analyses were performed using the PyMVPA software package (Hanke et al., 2009).

To determine the statistical significance of searchlight classification accuracies we used nonparametric randomization tests shuffling the entropy condition assignments (e.g., Etzel, 2015, 2017; Nastase et al., 2016). That is, for each permutation, the condition labels were randomly reassigned for all participants and the entire searchlight classification analysis (cross-validated across participants) was recomputed. For each searchlight analysis, 1,000 permutations were computed per searchlight, resulting in a distribution of searchlight maps under the null hypothesis of no systematic relationships among the condition labels. The actual searchlight classification accuracy was then compared against this null distribution to determine a p-value per searchlight. The permutation test respected the stratification of the data such that entropy labels were permuted within each participant, and for cross-classification permuted within each modality (nested within participants). When classifying highly-regular and random series (entropy levels 1 and 4), only labels 1 and 4 were permuted. When classifying extreme versus intermediate levels of entropy (quadratic profile), labels were permuted to ensure that the two extreme entropy levels were assigned the same label (“extreme” or “intermediate”) and that the intermediate levels were both assigned the other label.

We performed nonparametric cluster-level inference using a Monte Carlo procedure simulating clusters of random Gaussian noise (Forman et al., 1995). To constrain the spatial smoothness of the noise simulation, we computed residual searchlight accuracy maps by subtracting the average accuracy (across participants) from each participant’s accuracy (as in Linden, Oosterhof, Klein, & Downing, 2012, p. 630). The mean smoothness along the *x*-, *y*-, and *z*-axes of these residual accuracy maps was calculated using AFNI’s 3dFWHMx and supplied to AFNI’s 3dClustSim to estimate the extent of significant searchlight clusters occurring by chance. For the cross modal searchlight classification procedure, we report clusters that survived a voxel-level cluster-forming threshold of *p* < .05 (assessed using permutation tests) and a cluster-level threshold of *p* < .05, controlling for the family-wise error rate. For the within-modality searchlight classification, we use a slightly more conservative single-voxel cluster-forming threshold (*p* < .01; cluster-level correction, *p* < .05).

To relate the current study to prior reports of hippocampal sensitivity to statistical regularities (e.g., Bornstein & Daw, 2012; Covington et al., 2018; Schapiro et al., 2014; Turk-Browne et al., 2009), and because the searchlight approach is not particularly well-suited to subcortical structures, we additionally performed classification analyses within an anatomically defined hippocampal region of interest (ROI). For each participant, left and right hippocampal volumes were automatically segmented using FSL’s FIRST (Patenaude, Smith, Kennedy, & Jenkinson, 2011) and then normalized to MNI space following the same procedure described above for the whole brain. Voxels in MNI space assigned to hippocampus in 50% or more participants were included in the final hippocampal ROI. We then performed the classification analyses described above (i.e., cross-modal and within-modality, as well as linear and quadratic classification) within the hippocampus ROIs using leave-one-participant-out cross-validation. We analyzed the right and left hippocampal volumes separately. Significant classification within the hippocampus was assessed using the randomization test described above.

## 3. Results

### 3.1. Sensitivity to uncertainty in auditory, visual and audiovisual series

Using a multivariate searchlight analysis, we first evaluated two questions: *i*) whether local neural response patterns discriminate between highand low-uncertainty conditions (approximating a linear trend), and *ii*) whether response patterns discriminate the two intermediate levels of regularity from both the mostand least-regular conditions (approximating a quadratic trend). Responses discriminating high and low entropy are consistent with predictive processing, while responses discriminating intermediate and extreme levels of entropy may reflect processes modeling the complexity of the system generating the stimuli.

For the *auditory modality*, we identified two regions discriminating high and low entropy: left precentral gyrus (PCG) and right insula, with peak searchlight classification accuracies of 77–79%, cross-validated across participants (theoretical chance = 50% here and for all subsequent classification results). We found a much more extensive set of regions that discriminated the intermediate levels of auditory regularity from both high and low levels (a quadratic discrimination with respect to entropy levels; Figure 2). These included bilateral superior and middle temporal gyri (STG, MTG), right transverse temporal gyrus (TTG), occipital regions bilaterally, and the cerebellum, with peak two-way classification accuracies of 67–70% (see Table 1 for all significant cluster locations).

**Figure 2:**
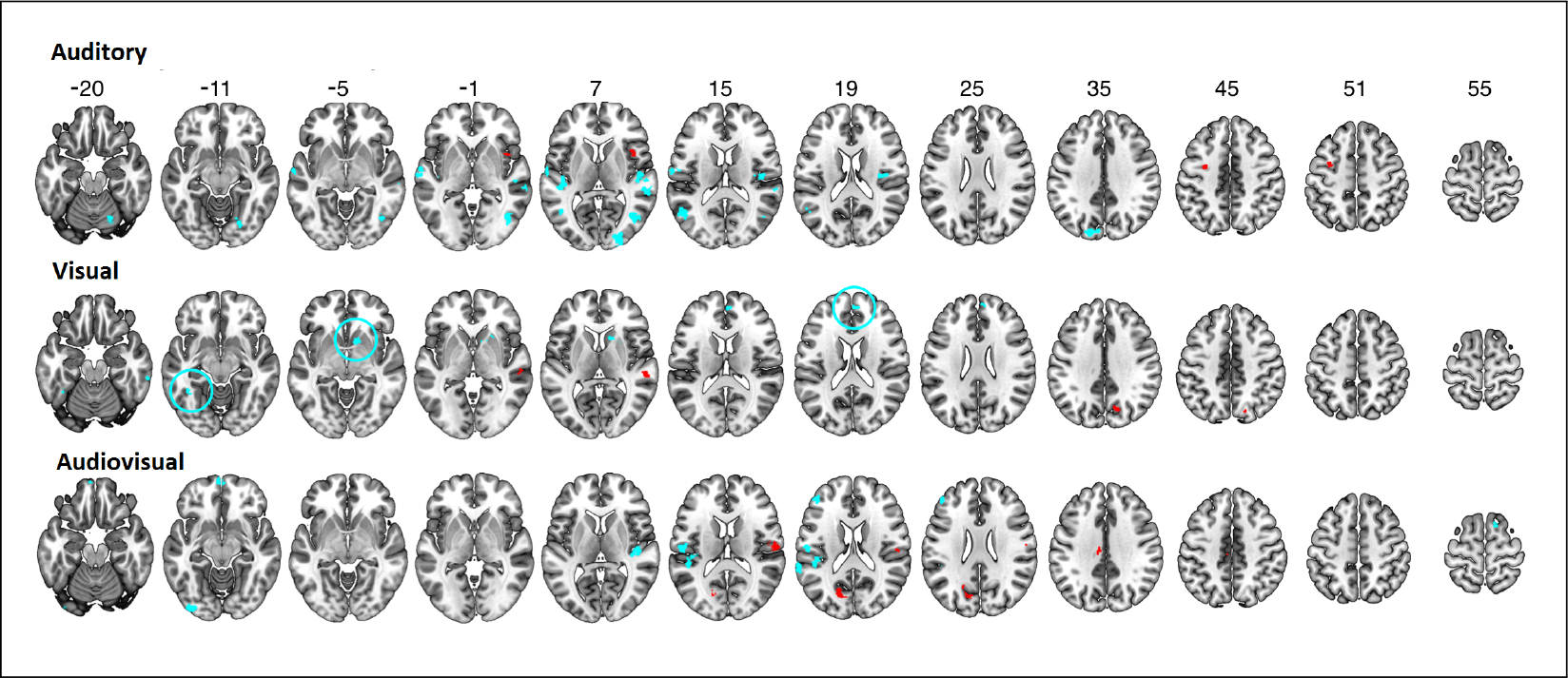
Neural sensitivity to uncertainty in the auditory, visual, and audiovisual series. Significant clusters of searchlights where response patterns reliably discriminated highly regular and random series are indicated in red. Significant clusters where response patterns discriminated the two intermediate levels of regularity from the two extreme levels (a quadratic discrimination) are plotted in cyan (cluster-level *p* < .05, corrected for multiple comparisons). Classifiers were tested using leave-one-participant-out cross-validation and statistically evaluated using permutation tests.

For the visual modality we identified four clusters discriminating high and low entropy series: left post-central gyrus (PoCG), right STG, right cuneus, and left rectal gyrus, with peak classification accuracies of 75-79%. We also identified several clusters discriminating intermediate levels of regularity from both the highest and lowest levels. The largest of the clusters was located in the right caudate, with additional clusters in the left fusiform gyrus, right inferior temporal gyrus, and right superior medial frontal gyrus. Peak classification accuracies in these significant clusters of searchlights ranged from 67% to 70%.

For the audiovisual stimuli, we identified three clusters that discriminated high and low entropy conditions: right PoCG, left superior occipital gyrus, and the left middle cingulate gyrus, with peak classification accuracies of 71-77%. Additionally, in several regions the classifier discriminated the intermediate and extreme entropy levels, including the left STG, right TTG, left PoCG, left cerebellum, and right supplementary motor area (SMA), with peak classification accuracies of 64-70%.

### 3.2. Cross-modal sensitivity to uncertainty

We used cross-modal searchlight classification to identify brain areas where response patterns discriminating levels of uncertainty generalized (i.e., were similar) across the auditory and visual modalities. Classifiers were trained to discriminate between levels of regularity in one sensory modality and then tested on input from the other modality using leave-one-participantout cross-validation. Cross-modal classification of high and low entropy series revealed significant clusters of searchlights in left orbitofrontal cortex (OFC), left MTG, and right cerebellum (see Figure 3, red clusters; cluster-level p < .05, corrected for multiple comparisons). These significant clusters had peak classification accuracies of 66-70% cross-validated across participants (theoretical chance = 50%). Interestingly, cross-modal classification discriminating intermediate and extreme levels of entropy (analogous to a U-shaped discrimination) was extensive (Figure 3, cyan clusters), implicating the right inferior frontal gyrus, SMA and SMG bilaterally, left PCG and MFG, left cerebellum, left superior parietal lobule, left inferior occipital and fusiform gyri, and the right insula (cluster-level p < .05, corrected for multiple comparisons; see Table 2 for all significant clusters and Supplementary Movie for 3D rendering). These areas exhibited peak two-way classification accuracies of 61-64%.

**Figure 3:**
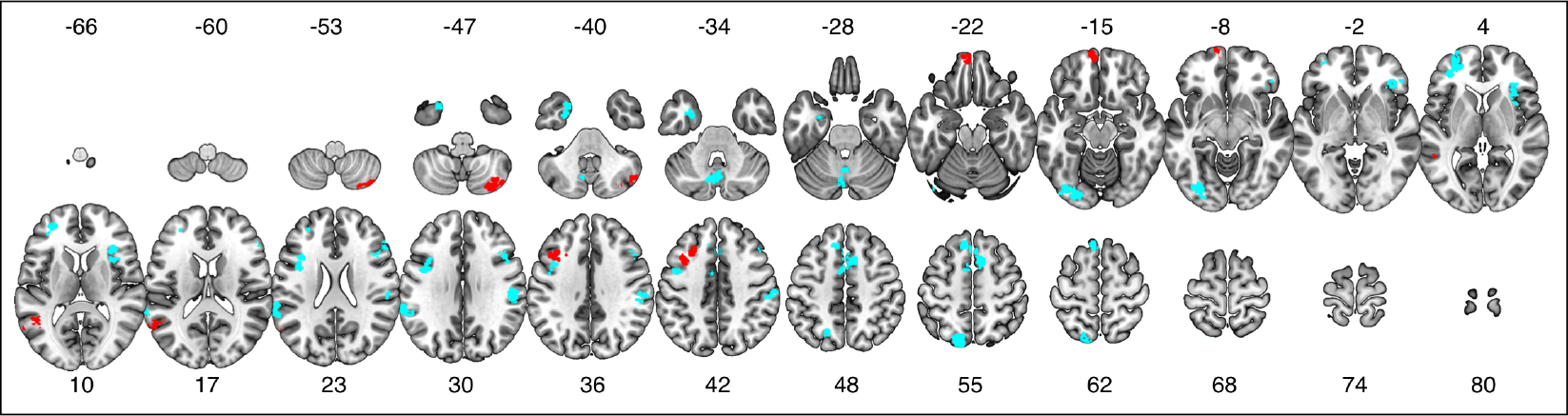
Cross-modal searchlight classification of input uncertainty. The searchlight analysis identified brain regions where classifiers trained to discriminate levels of regularity in one modality (e.g., auditory inputs) could successfully classify levels of regularity in the other modality (e.g., visual inputs, and vice versa). Significant clusters of searchlights where response patterns for highly regular and random series could be reliably classified across the auditory and visual modalities are indicated in red. Significant clusters of searchlights with reliable cross-modal classification of intermediate and extreme levels of regularity are plotted in cyan. Classifiers were tested on the left-out modality using leave-one-participant-out cross-validation, and statistically evaluated using permutation tests. Searchlight results are statistically significant at cluster-level p < .05, corrected for multiple comparisons.

### 3.3. Hippocampal analysis

Given prior studies implicating the hippocampus in associative learning, statistical learning, and sensitivity to uncertainty more generally (Strange et al., 2005; Harrison et al., 2006; Turk-Browne et al., 2009; Reddy et al., 2015), we conducted multivariate analyses analogous to those reported for searchlights above within anatomically-defined left and right hippocampus ROIs. Given their potentially differential roles in contextual integration (see Hartzell, Tobia, Davis, Cashdollar, & Hasson, 2015), we separately analyzed response patterns in the left and right hippocampus. As in the previous analyses, we tested for responses discriminating both high and low entropy levels, and intermediate and extreme entropy levels for auditory, visual, and audiovisual stimuli (thus six tests per left and right hippocampus, 12 total). In addition, we evaluated cross-modal linear and quadratic classification analyses (two tests per left and right hippocampus, four total). Due to the exploratory nature of the analysis we did not control for family-wise error over the 16 tests.

The analysis yielded two suggestive findings. For the right hippocampus, response patterns discriminated highly regular and random audiovisual series with 71% accuracy (theoretical chance = 50%, p = .003, permutation test, uncorrected). In the left hippocampus, cross-modal classification of highly regular and random series reached 62% accuracy (p = .004, permutation test, uncorrected). Apart from these two instances, all other tests yielded accuracies very close to or below chance level.

## 4. Discussion

Our main aims were to determine whether it is possible to identify monotonic and non-monotonic neural sensitivity to uncertainty, and whether these neural signatures generalize across sensory modalities. Our main findings are as follows. First, multivariate pattern classification analysis proved highly sensitive, suggesting that neural response patterns contain information differentiating levels of uncertainty in short input series. Response patterns not only discriminated between highly regular and random series, which is to be expected from the existing literature (e.g., Cunillera et al., 2009; McNealy et al., 2006), but in some areas also discriminated the two intermediate levels of uncertainty from the two extremes. This latter finding is consistent with the view (e.g., Nastase et al., 2014; Kidd et al., 2012; Hasson, 2017) that sensitivity to input statistics may comprise computations for which neural activity does not scale monotonically with uncertainty. The resulting network of brain regions includes perisylvian areas implicated in prior work (e.g., Tobia et al., 2012) and overlaps with areas characterized by intermediate-scale temporal receptive windows (Hasson et al., 2008; Lerner, Honey, Silbert, & Hasson, 2011). Uncertainty in audiovisual stimuli engaged some systems recruited in the auditory condition (e.g., superior temporal cortex), but also recruited prefrontal systems.

Equally important, when probing for supra-modal sensitivity to uncertainty, we identified a number of regions where it was possible to decode the level of uncertainty in one modality using a classifier trained discriminate levels of uncertainty in the other modality. Interestingly, cross-modal responses differentiating intermediate from extreme levels of entropy comprised a fairly widespread network, whereas relatively few areas differentiated high and low levels of uncertainty in a cross-modal fashion.

### 4.1. Cross-modal and non-monotonic sensitivity to uncertainty

Few studies have directly examined whether there are neurobiological systems that track the level of regularity in sensory inputs in a supra-modal fashion; that is, independently of sensory modality. Two studies by Schubotz et al. compared processing of regularity in either auditory and visual series (Schubotz & von Cramon, 2002) or abstract series and motor actions (Schubotz & von Cramon, 2004), reporting adjacent and overlapping activity patterns in premotor cortex. However, both studies required participants to deliberately predict future events, which makes it unclear whether the results are a result of endogenous processing or explicit executive demands. In our own prior work (Nastase et al., 2014), we identified areas sensitive to regularity for either auditory or visual inputs, but found no area that was generally sensitive in both modalities. As noted in the Introduction, conjunctions of unisensory response maps provide only weak evidence for abstract, supra-modal computations (e.g., Peelen & Downing, 2006). Rather, in the current study we used cross-modal classification, which provides a more robust test of representational content shared across modalities (Man et al., 2012; Kaplan et al., 2015).

The cross-modal searchlight classification analysis identified an extensive supra-modal network of regions, some discriminating highly regular from random inputs (the typical contrast in univariate studies of uncertainty), and others differentiating the intermediate and extreme conditions in a quadratic fashion. Cross-modal regions discriminating the most regular from random inputs were limited to the left posterior middle temporal gyrus, the left premotor cortex/middle frontal gyrus, and left orbitofrontal cortex. The posterior lateral temporal cortex is multisensory, receiving input from both auditory and visual association cortices (Barnes & Pandya, 1992; Beauchamp, Argall, Bodurka, Duyn, & Martin, 2004). It may be that multi-modal temporal areas sensitive to regularity in the environment are recruited similarly across modalities, and prior work has shown that this area tracks regularity in visual series (Bischoff-Grethe, Proper, Mao, Daniels, & Berns, 2000). Cross-modal sensitivity to uncertainty in premotor cortex is consistent with prior findings (Schubotz & von Cramon, 2002) documenting its sensitivity to the complexity of auditory and visual series, though in non-overlapping areas. Meyniel and Dehaene (2017) have also linked this region to tracking confidence and uncertainty in simple auditory and visual series.

Cross-modal sensitivity to uncertainty observed in orbitofrontal cortex is consistent with recent theories implicating this region, and limbic cortices more generally, as a source of predictive feedback signals conveyed to lower-level perceptual areas (Chanes & Barrett, 2016; Trapp & Bar, 2015). As the highly regular and random series differ in the extent to which they license predictions, observed differences in the response topographies in these areas may reflect the operation of predictive processes. Finally, in a post-hoc analysis, we discovered that response patterns in the right hippocampus differentiate highly structured and random audiovisual series, while the left hippocampus differentiates structured and random series across modalities. These suggestive results support previous work pointing to associative learning in the hippocampus in the context of implicit learning (Turk-Browne et al., 2009, Reddy et al., 2015). In future work, cross-modal classification may prove useful in testing whether the hippocampus encodes statistical regularities in a supra-modal fashion.

Cross-modal responses discriminating the intermediate and extreme levels of regularity were surprisingly prevalent (seen in the spatial extent of cyan clusters in Figure 3). On the left, these were found in the middle frontal gyrus, temporal pole, lateral occipital cortex, intraparietal sulcus, and auditory association cortex. On the right, significant clusters were identified in the supramarginal gyrus and inferior frontal gyrus. Immediately rostral to the premotor cluster that discriminated high and low entropy conditions cross-modally, we identified another cross-modal cluster that discriminated intermediate from extreme entropy levels. Schubotz et al. (2002) had suggested that different areas in premotor cortex are involved in prediction of auditory and visual stimuli, such that prediction of auditory sequences utilizes premotor areas involved in verbal articulation, and prediction of visual sequences utilizes areas involved in hand movement. Expanding on this idea, our findings suggest a finer-grained, common substrate for the representation of uncertainty across modalities in premotor cortex. Quadratic entropy discrimination was also found in the dorsomedial prefrontal cortex bilaterally. We have previously documented an analogous type of non-monotonic, U-shaped response profile to regularity in auditory series when considering short (10 s) epochs (Tobia, Iacovella, Davis, & Hasson, 2012), as well as U-shaped whole-brain connectivity profiles for the anterior cingulate cortex during long periods of auditory stimulation (Nastase et al., 2015).

These quadratic, non-monotonic response profiles are compatible with several types of computational accounts, as we have previously discussed in detail (Nastase et al., 2015; Hasson, 2017). In brief, they may be indicative of systems that do not explicitly code for statistical predictability or regularity per se, but instead are involved in modeling the system generating the sensory input. This modeling process may be outwardly reflected in an apparent U-shaped response profile because such model descriptions are simpler to construct for environmental systems that generate highly regular or random inputs than for systems that generate inputs with intermediate levels of regularity. Another possibility is that these brain areas subserve prediction of multiple future transitions (e.g., two stimuli into the future, *t* + 1, *t* + 2), and are sensitive to *predictive information rate* (PIR): the degree to which the stimulus at time *t* + 1 impacts the observers certainty regarding the stimulus expected at *t* + 2. Computational work has shown that PIR is maximal for series with intermediate levels of disorder, but lowest for both very regular and random series (Abdallah & Plumbley, 2009). A third possibility is that these brain areas are involved in the generation of predictions and sensitive to prediction error, but only so long as predictions are licensed by the input. This might be reflected in gradually increasing activity as disorder increases within a reasonable bound, but with a subsequent decline for the random condition, where no predictions are licensed. Thus, both the highly ordered and random condition would be accompanied by low prediction errors. We note, however, that the latter interpretation may be the least plausible for the relatively brief 10 s series presented here, because for such short random series it may be quite difficult to establish evidence that prediction is not licensed, particularly given the tendency to perceive streaks in completely random inputs (Huettel et al., 2002).

Areas encoding audiovisual entropy were largely non-overlapping with areas encoding entropy in unisensory visual and auditory series, which is consistent with prior work (Andric, Davis, & Hasson, 2017). This relatively modest overlap is also consistent with behavioral work suggesting that multisensory regularities are learned independently of regularities conveyed via their unisensory constituents (Seitz, Kim, van Wassenhove, & Shams, 2007). In the current experiment, the auditory and visual channels in the audiovisual condition provided redundant statistical information (mutual information was maximal). This may have produced a more efficient encoding of the series tokens themselves, in this way affording greater sensitivity to audiovisual regularities.

Although participants performed an orthogonal cover task, we cannot rule out the possibility that implicit attentional allocation may have co-varied with the entropy manipulation. Attention and prediction are related constructs and often conflated experimental work (Summerfield & Egner, 2009), where statistical regularities licensing expectations are often used to guide attention (Posner, Snyder, & Davidson, 1980; Zhao, Al-Aidroos, & Turk-Browne, 2013). Recent work on the interaction of these processes has met with mixed results (Jiang, Summerfield, & Egner, 2013; Kok et al., 2012) and our experiment was not designed to adjudicate between these processes. Note that our procedure for normalizing response patterns may be robust to simple attentional effects (by mean-centering each searchlight), but does not necessarily rule out more complex attentional effects (Jehee, Brady, & Tong, 2011; Nastase et al., 2017).

Our findings point to the importance of examining non-monotonic responses to predictability and uncertainty when studying brain systems sensitive to input statistics, as such responses may be as prevalent as the morestudied monotonic response profiles. More generally, they demonstrate the utility and importance of using cross-modal classification for drawing conclusions about supra-modal computations underlying statistical learning. While work to date, including our own, has largely failed to identify supra-modal systems sensitive to sequential structure, suggesting that sensory cortices support these computations (e.g., Dehaene et al., 2015; Nastase et al., 2014; Schubotz & von Cramon, 2002), this conclusion may rely in part on analytic limitations. The cross-classification approach used here suggests that widespread association cortices are sensitive to structure in sequential stimuli across sensory modalities.

### 4.2. Methodological considerations

Multivariate approaches provide specific insights into distributed neural representation, with prior studies suggesting that searchlight pattern analyses are both more sensitive and more opportunistic in exploiting potential confounding variables (Coutanche, 2013; Davis et al., 2014; Jimura & Poldrack, 2012). As such, several considerations should be discussed when interpreting the current findings. First, the searchlight approach provides coarse localization, as each searchlight aggregates information over numerous (i.e., over 100) voxels and overlaps with numerous neighboring searchlights. Furthermore, to better approximate other correlation-based classification analyses (e.g., Haxby et al., 2001), we normalized (i.e., Z-scored) response patterns across voxels within each searchlight prior to classification (Misaki et al., 2010; Nastase et al., 2016). This procedure effectively removes any searchlightaverage differences in response magnitude between experimental conditions. The classifier therefore operates solely on distributed response topographies of relative activity, and cannot capitalize on general differences in overall response magnitudes across conditions within a given searchlight. Abandoning this normalization scheme and allowing the classifier to also utilize regional-average response magnitudes would likely more closely approximate a conventional univariate analysis.

Davis et al. (2014) argued that multivariate analyses may appear to offer greater sensitivity than univariate analyses because they exploit idiosyncratic within-participant response variability that is typically discarded at the group level in univariate analyses. However, this concern holds primarily for within-participant classification analyses where the result of classification (e.g., classification accuracy, cross-validated across runs within a participant) is then aggregated in a group level statistical analysis. In contrast, here we used leave-one-*participant*-out cross-validation, which limits classifiers to voxel-level response variability that is consistent across participants. Allefeld and colleagues (2016, pp. 382-383) have demonstrated that performing second-level statistical tests on participant-level classification accuracies (which are typically distributed asymmetrically above chance) does not properly perform population-level inference (effectively testing only the global null hypothesis that there is no effect for any participant). When separate classifiers are trained per participant, accuracies may result from idiosyncratic patterns that distinguish conditions in one individual but do not generalize to other individuals. Allefeld and colleagues suggest that performing crossvalidation across participants, on the other hand, effectively provides rigorous population inference. However, this approach comes with a potential cost. Specifically, leave-one-participant-out cross-validation requires that response patterns are spatially registered across participants (within the radius of a searchlight). Although all participants’ data were spatially normalized to the MNI template prior to classification, anatomical alignment cannot in principle perfectly align fine-grained functional topographies and yields differentially effective alignment across the brain (Guntupalli et al., 2016; Haxby et al., 2011). While our results add to previous work in demonstrating that cross-participant classification is feasible (e.g., Mourao-Miranda et al., 2005), due to the imperfect registration of functional topographies, classification may rely on relatively coarse-grained response topographies differentiating levels of uncertainty.

### 4.3. Summary

To date, relatively modest progress has been made in developing neurobiological accounts of uncertainty that extend beyond explanations of monotonic responses in single modalities. The current study informs current neurobiological models of neural sensitivity to statistical regularities in two ways. 21 667 668 669 670 671 672 We identified neural systems that encode information about statistical regularities in a supra-modal manner, as evidenced by cross-modal multivariate classification. In addition, our findings emphasize that the human brain responds to uncertainty both monotonically and non-monotonically, suggesting that some brain regions track uncertainty per se, while others code for structural features of the systems generating the sensory input.

